# Cryo-Electron Microscopy Visualization of a Large Insertion in the 5S ribosomal RNA of the Extremely Halophilic Archaeon *Halococcus morrhuae*

**DOI:** 10.1101/2020.05.05.079889

**Authors:** Madhan R Tirumalai, Jason T Kaelber, Donghyun R Park, Quyen Tran, George E Fox

**Affiliations:** Dept of Biology and Biochemistry, University of Houston, Houston, TX 77204-5001 USA; National Center for Macromolecular Imaging, Baylor College of Medicine, Houston, TX 77030, USA

**Keywords:** Expansion sequences, Insertion sequences, accretion model, archaea, ribosomal RNA

## Abstract

The extreme halophile *Halococcus morrhuae* (ATCC® 17082) contains a 108-nucleotide insertion in its 5S rRNA. Large rRNA expansions in Archaea are rare. This one almost doubles the length of the 5S rRNA. In order to understand how such an insertion is accommodated in the ribosome, we obtained a cryo-electron microscopy reconstruction of the native large subunit at subnanometer resolution. The insertion site forms a four-way junction that fully preserves the canonical 5S rRNA structure. Moving away from the junction site, the inserted region is conformationally flexible and does not pack tightly against the large subunit.

## INTRODUCTION

Single molecule fluorescence resonance energy transfer (smFRET), crystallographic and cryo-electron microscopy (cryo-EM) techniques have been used to obtain high resolution structures of ribosomes in various stages of its motions during translation as well as its various components (Frank and Agrawal 2000; Valle et al. 2003; Gagnon et al. 2016; Shebl et al. 2016; Wasserman et al. 2016; Zhou et al. 2020). These studies have been instrumental in our understanding of ribosomal evolution.

Ribosomal diversity has emerged largely by stepwise addition of insertion sequences that “grow” the ribosome outward, from the common core to the large ribosomes of extant complex metazoans (Bokov and Steinberg 2009; Petrov et al. 2015). In some instances, it has been possible to recognize that what is now a common part of the RNA likely began as an insertion (Yokoyama and Suzuki 2008; Hsiao et al. 2013; Petrov et al. 2015; Gomez Ramos et al. 2017). Such rRNA insertions have been used to deduce the relative age of various regions in those RNAs (Petrov et al. 2015). Insertions typically are accommodated in the ribosome by forming a three or four-way junction with negligible perturbation of the parental helix (Petrov et al. 2014).

5S rRNA as an integral part of the ribosomal large subunit LSU may function as a link between the peptidyl transferase center and GTPase center of the ribosome via loop D (Dontsova et al. 1994; Dokudovskaya et al. 1996; Sergiev et al. 2000; Smith et al. 2001; Dontsova and Dinman 2005; Kouvela et al. 2007) as well as through its interacting proteins uL5, uL18 and L25 (Christiansen and Garrett 1986; Lu and Steitz 2000; Yusupov et al. 2001; Smirnov et al. 2008)..

Of special interest here is the 5S rRNA isolated from *Halococcus morrhuae* (ATCC® 17082), which contains a large 108 nucleotide insertion (Luehrsen et al. 1981) that almost doubles the size of the RNA (Fig. 1). The insertion is found between *H. morrhuae* residues 104 and 105 (Luehrsen et al. 1981), which corresponds to residues 108 and 109 in the universal bacterial numbering system (Fox 1985). All members of the genus *Halococcus* that have been examined bear this insertion (Nicholson 1982; Stan-Lotter et al. 1999). Although insertion/expansion sequences of varying size are frequently seen in the large rRNAs of eukaryotes (Gerbi 1996; Taylor et al. 2009), similar large insertions have typically not been described in archaea (Petrov et al. 2014). Exceptions occur in the Asgard-group archaea that may be related to eukaryotes (Penev et al. 2020). In bacteria, there has been at least one notable instance, wherein an insertion known as the “steeple” (causing a small-subunit-dependent conformational change) has been described in *Mycobacterium smegmatis* (Shasmal and Sengupta 2012; Yang et al. 2017). The fact that the *H. morrhuae* insert is almost as large as its parent rRNA raises the question of whether the presence of the insert may impact the canonical 5S structure. In order to address this issue, cryo-EM was used to characterize the 3D structure of the native large subunit of *H. morrhuae* (ATCC® 10782).

**FIGURE 1.**
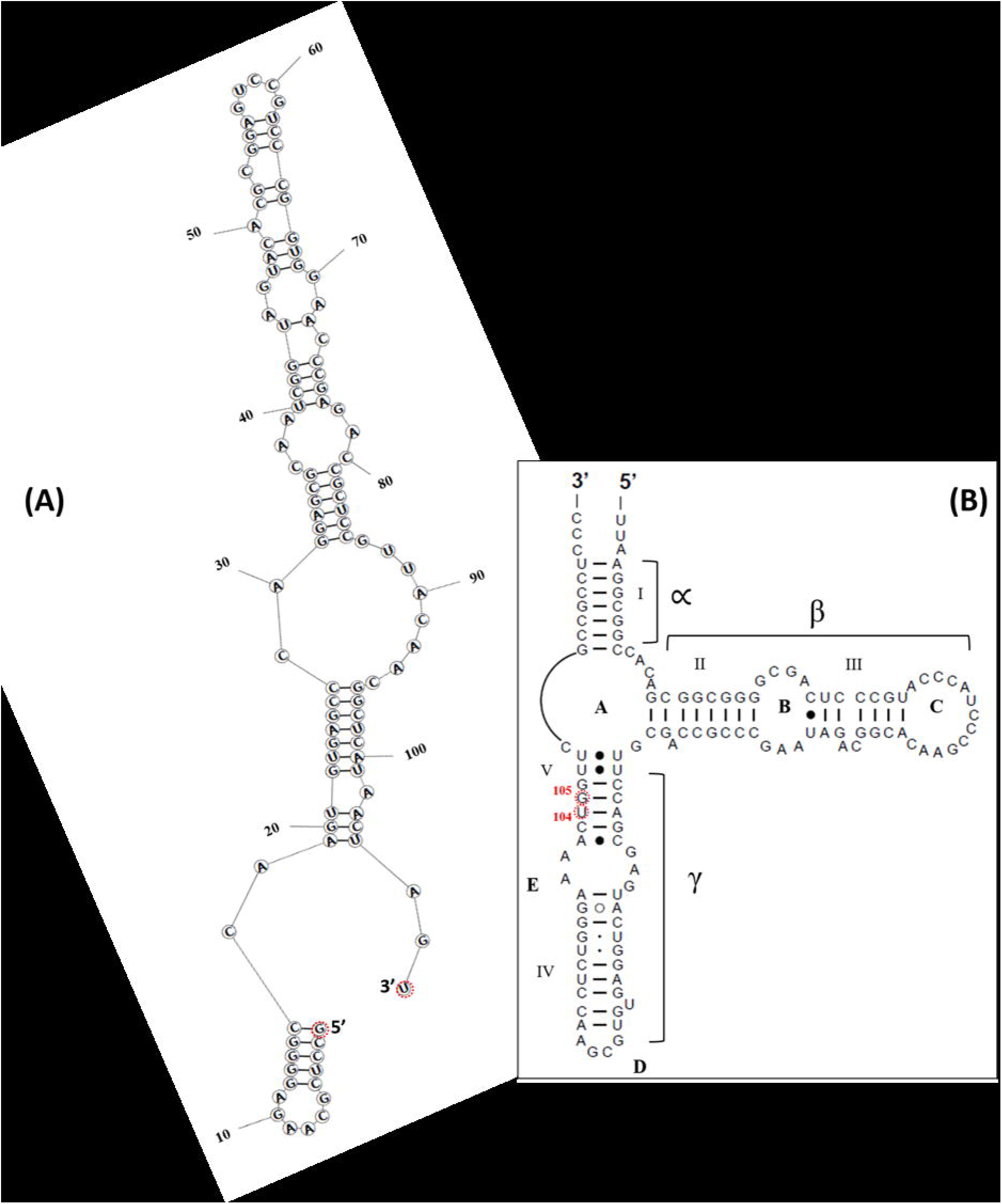
A schematic diagram showing (A) a possible secondary structure of the insert as predicted by mFold (Zuker 2003) and (B) the usual 5S rRNA secondary structure (obtained from RNAcentral (http://www.rna.ccbb.utexas.edu) (Cannone et al. 2002; RNAcentral Consortium 2019). The insert is between positions U104 and G105 as per *Halococcus morrhuae* 5S rRNA numbering. The equivalent positions in *Haloarcula marismortui* are C108 and G109. What would be the 5’ and 3’ends of the insert if it were an independent RNA are indicated in (A).

## RESULTS AND DISCUSSION

Secondary structure prediction suggests several alternative structures for the insert. The insert structure with the most negative delta G (−44.8 kcal/mol) (delta G values are inversely proportional to the stability of the secondary structure) (Fig. 1A) suggests that the insert alone would form an elongated structure with five helical regions each consisting of four or more standard base pairs. From a transcriptional perspective, synthesis would progress from the 5’end of helix 1, which would then be completed before the other helices are even started. This would result in a small structural domain consisting of helix 1 alone and a large domain encompassing the rest of the insert. The predicted helices in the insert are consistently separated by bulge loops that likely allow changes in orientation such that the RNA can wrap around the 50S subunit.

We purified large ribosomal subunits from *H. morrhuae* ATCC® 17082 and imaged them by cryo-EM. This allowed us to obtain a 3D model of the 5S rRNA in the context of the intact large subunit. In pilot cryo-EM experiments on ribosomes (extracted from *H. morrhuae* ATCC® 17082) at low salt concentrations, the quality was poor, likely due to their lack of stability at these concentrations (Ring and Eichler 2004). Therefore, ribosomes were maintained in the high-salt Buffer A until vitrification. The salt concentration was then lowered with an on-grid washing procedure. Initial reconstructions to 12Å revealed density protruding from the expected site of the 5S rRNA. This protrusion is not present in known LSU structures such as that of *Methanothermobacter thermautotrophicum* (Fig. 2D) (PDB ID 4ADX) and *H. marismortui* (Klein et al. 2001; Greber et al. 2012). Docking known 5S structures into the map showed that the site of the protruding density matches, to the nucleotide, the expected site of the insertion assuming no insertion-induced changes of the 5S rRNA (Fig. 1; Fig. 2C-D) The protruding density fades and broadens as the distance from the core increases, which could indicate flexibility of the looped-out region.

**FIGURE 2.**
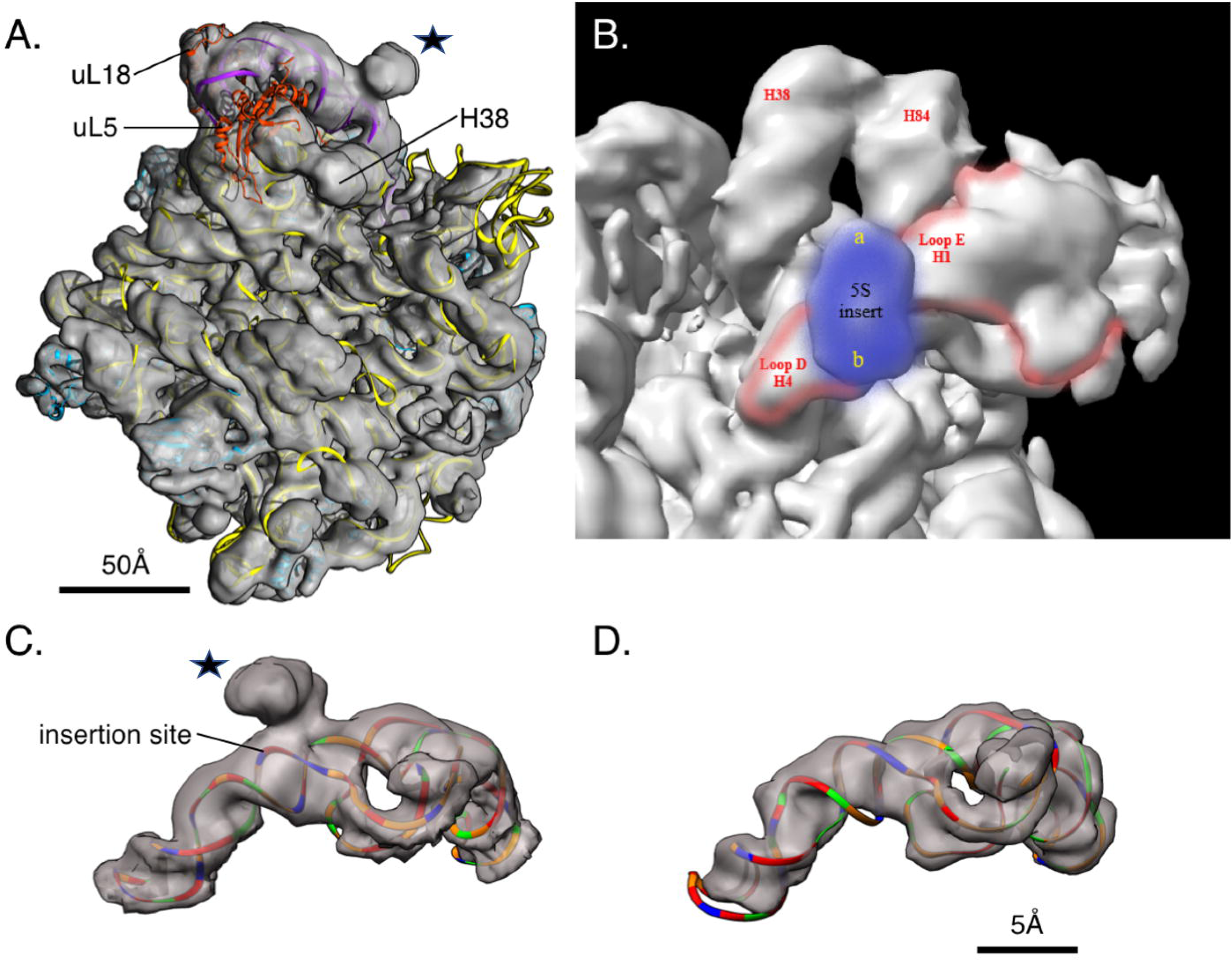
(A) Cryo-EM reconstruction of the large subunit of *H. morrhuae*. The large subunit of *Haloarcula marismotui* (PDB: 1NJI) (Hansen et al. 2003) is rigidly docked in the density; 5S rRNA (purple), 23S rRNA (yellow), uL5 and uL18 (orange-red), and selected ribosomal other proteins (cyan) are shown. The insert is marked by ★. (B) Top view of the 50S subunit. Features surrounding the 5S rRNA extension (blue) are labeled and the canonical 5S rRNA is outlined in red. (C) The section of the map corresponding to the 5S rRNA is cropped out for visualization. The *Haloarcula marismortui* 5S rRNA was mutated to match the sequence of the *Halococcus morrhuae* 5S, except for the insert whose secondary structure is not known, and rigidly docked inside the density. Coloring is by nucleotide: adenines, green; cytosines, orange; guanines, red; and uracils, blue. The insert is marked by ★. (D) The section of the map corresponding to the 5S rRNA is cropped out of the published, 6.6Å *Methanothermobacter thermautotrophicus* cryoEM map and associated atomic model (EMD-2012, PDB 4ADX) (Greber et al. 2012) and colored as in (C). This and all other homolog maps lack the protruding lump of density seen in (C).

Using freshly-prepared LSUs the imaging was repeated at higher magnification and with a larger dataset of over 100,000 particles. This data was reconstructed to a nominal resolution of 6.4Å (EMD-21670) (Fig. 2A-D). RNA and protein secondary structural elements are clearly seen in the interior of the particle (Fig. 2A). The density seen for the insertion still projects off of base 108 of the 5S rRNA (Fig. 2C), but it is poorly resolved and only large enough to accommodate around 40 of the 108 inserted nucleotides, as a continuous double helix. The density suggests a helix extending at 120° and 210° relative to the parent helix in a four-way junction (Fig. 2B). The junction resembles the specificity domain four-way junction of RNase P (Krasilnikov et al. 2003), which is the type specimen of four-way junction family “H”(Laing et al. 2011). The fact that the bulge does not accommodate all of the insert suggests that the insert may “wave” like a flag in the wind. An abrupt cutoff is not seen but rather a gradual broadening and fading of the density.

At low isosurface thresholds, some density can be seen stretching from the insertion site to helix 38, the A site loop near *E. coli* positions 970-990 at the subunit interface. Helix 38 makes a lateral contact with helix 84 in the 23S rRNA (Fig. 2B). Helix 38 houses the A-site finger that aids in positioning the tRNA in the A-site (Stark et al. 1997; Nissen et al. 2001). The faint density does not conflict with the locations of the nearby ribosomal proteins uL18 and uL5 (Fig. 2A). uL5 density is fainter than uL18 density, potentially indicating partial occupancy in this dataset. However, from these maps we are not able to determine whether the faint density stretching from the insertion site to helix 38 represents genuine, flexible 5S rRNA density or noise.

No other post-LUCA ribosomal insertion has almost doubled the size of a ribosomal RNA. Yet, even this large insertion obeys the general rules that have been proposed for ribosomal accretion (Petrov et al. 2014) in that it leaves a typical insertion fingerprint and causes negligible rearrangement of nucleotides in the parent helix. The apparent flexibility of the inserted nucleotides is atypical. The observed flexibility could be an intrinsic feature of this rRNA insertion reflecting its age or function, or the insertion might assume a homogeneous conformation in the presence of a binding partner (Shasmal and Sengupta 2012; Hentschel et al. 2017; Yang et al. 2017). The flexibility of the inserted region within the halococcal 5S rRNA and its evolutionary history warrant future study.

Aside from the 5S rRNA, comparing the *H. morrhuae* large subunit map obtained herein to the crystal structures of *H. marismortui* crystal structure (Evers et al. 1994; Penczek et al. 1999; Gabdulkhakov et al. 2013) reveal minor differences at the subunit exterior. In this earlier map, but not in crystals, the end of helix 38 is seen (Fig. 2A). On the opposite side of the 5S insertion site is helix 25. Whereas in the crystal structure this helix wraps against the ribosome, in solution it extends outward. The flipped bases such as 578-580 in the crystal structure are likely crystal packing artifacts. The structures also differ at *E. coli* positions 2865-2890 (Helix 101) where the RNA continues in a different direction than in the crystal structure also likely an effect of crystal packing.

## MATERIALS AND METHODS

### Computational analysis

A prediction of the insert’s likely secondary structure was made using mfold (Zuker 2003) and RNAstructure (Reuter and Mathews 2010). The 5S rRNA secondary structure without the insert was obtained from RNAcentral (http://www.rna.ccbb.utexas.edu) (Cannone et al. 2002; RNAcentral Consortium 2019).

### Medium, culture / strain and maintenance

*Halococcus morrhuae* (Farlow) Kocur and Hodgkiss (ATCC^®^ 17082™) was obtained from the ATCC and maintained on 25% NaCl media containing casamino acids, and other salts (Nicholson 1982).

### Cell growth and lysis for extraction of the ribosomes

*Halococcus morrhuae* cells were grown on a large-scale using a minimum of one liter of culture broth, at 37°C, 200 rpm. The cells were harvested during mid-log phase, suspended in buffer A (3.4 M KCI, 60 mM Mg (OAc), 30 mM Tris-HC1, 7 mM 2-mercaptoethanol, pH 7.6 (Sanchez et al. 1990). Suspended cells were passed through a French press multiple times. The cell lysates were treated with RNAse free DNAse (Promega Corp) for 30 min, following which the lysate was processed through ultra-centrifugation as described (Sanchez et al. 1990; Spedding 1990). As a first step, the cell lysates were centrifuged at 30000×*g* for 30 min to remove the debris in a SW28 rotor using a Beckman Coulter Ultracentrifuge. The upper two-thirds of the supernatant was collected and centrifuged in a SW28 rotor, 20,000 rpm for 17hrs. In one previous study involving ribosome isolation from *H. marismortui*, the red gelatinous material, presumably containing the pigments, that was found atop the ribosome pellet, was removed physically (Shevack et al. 1985). However, in the case of *H. morrhuae*, the ribosome pellet was found to be mixed with pigments, which could be an impediment to cryo-EM reconstruction. Ribosomes have been shown to be precipitated by the addition of acetone or ethanol (Suh and Limbach 2004). The ribosomes were separated from the pigments, by treating the ribosome pellet with acetone (100%) in a 50ml tube. The tube was intermittently inverted for 15-20 min to allow the separation of the pigments into acetone. The acetone phase was then carefully removed using a sterile pipette. The ribosomal pellet was air-dried, and suspended in buffer A. To obtain the ribosomal subunits, ribosomes in buffer A were diluted approximately 10-fold with dissociation buffer (2.7 M KCI). Next they were separated for 15 hours by zonal centrifugation using a linear, 6-36% (w/v), sucrose gradient in a SWTi 55 rotor, at 22,500 rpm (100,000×*g*) (Sanchez et al. 1990). The subunits were then pooled separately from the fractions, washed 2-3 times with buffer A to remove the sucrose and finally stored in buffer A at −80°C.

### Cryo-electron microscopy

Ribosomes were vitrified and imaged by cryo-EM. Preliminary images in Buffer A presented low contrast, presumably attributable to the high salt concentration. We therefore applied an on-grid washing procedure. Quantifoil R2/1 grids were plasma cleaned for 5 seconds using a Solarus plasma cleaner (Gatan). In the humidity-controlled chamber of the Vitrobot Mark IV, 2μL sample was applied to the grid, then 6μL of distilled water were applied to the reverse face of the grid and immediately blotted. The droplet of water on the reverse face was observed to travel through the grid and mainly pool on the plasma-cleaned face. A typical blot time for grids used in this study was 3 seconds.

Micrographs were collected using a JEOL JEM-3200FSC electron microscope with Gatan K2 Summit direct electron detector. The final dataset was imaged at 1.97Å/pixel, nominally 20,000× magnification, with a total exposure of 26e-/Å^2^ over 8 seconds. 1000 were collected over the course of two imaging sessions, of which 795 were accepted for processing. Subnanometer reconstructions were obtained using both Relion-2 and cryoSPARC v1. Data was traceably archived using the EMEN2 object-oriented database (Rees et al., 2013) until processing methods improved. After further developments in cryo-EM software, the dataset was reprocessed. Relion-3.1 was used for motion correction (Zivanov et al., 2019), CTF fitting, and particle extraction. crYOLO (Wagner et al., 2019) was used for neural-net-based particle picking. 140,444 particles were picked. The subset of particles exhibiting neither aggregation nor self-self-interaction was manually extracted for reconstruction. 2D classification to a subset of 99,337 particles, *ab initio* reconstruction and non-uniform refinement were performed in cryoSPARC v2 (Punjani et al., 2017). More stringent classification to a 69,042-particle subset did not improve the structure, nor did particle-subtracted refinement of the 5S rRNA alone.

For structure interpretation, known structures from the related halophilic archaeon *H. marismortui* (Ban et al. 1998; Penczek et al. 1999; Gabdulkhakov et al. 2013) and the relatively distant methanogenic archaeon *Methanothermobacter thermautotrophicus* (Greber et al. 2012) were docked onto the electron density of the *H. morrhuae* ribosome particles. Maps were rendered using UCSF Chimera or USCF Chimera X (Goddard et al. 2018). Segmentation of the map to isolate the 5S region (Fig. 2C-D) was performed in Chimera by using the atomic models of archaeal ribosomes and applying the “Split Map” command.

## SUPPLEMENTAL MATERIAL

Supplemental material is available for this article.

## ACCESSIONS

The cryoEM reconstruction of the *Halococcus morrhuae* (ATCC® 17082) large subunit is deposited as EMD-21670.

## ACKNOWLEDGEMENTS

This work was supported in part by NASA Exobiology Grants NNX14AK36G, NNX14AK16G and NASA Contract 80NSSC18K1139 under the Center for Origin of Life, Georgia Institute of Technology to GEF and NIH grant P41GM103832 to Wah Chiu. We thank Jessica Chin and Joanita Jakana for technical support.

## Author contributions

GEF conceived the work; GEF, MRT and JTK planned the work; MRT maintained the culture, grew the cells, extracted, and purified the ribosomes; JTK and DRP performed cryo-EM experiments; MRT and JTK constructed figures; MRT, JTK, QT and GEF analyzed the structure; all authors co-wrote the paper.

